# Intermittent Starvation Extends the Functional Lifetime of Primary Human Hepatocyte Cultures

**DOI:** 10.1101/818203

**Authors:** Matthew D. Davidson, Salman R. Khetani

## Abstract

Primary human hepatocyte (PHH) cultures have become indispensable to mitigate the risk of adverse drug reactions in human patients. In contrast to de-differentiating monocultures, co-culture with non-parenchymal cells maintains PHH functions for 2-4 weeks. However, since the functional lifespan of PHHs *in vivo* is 200-400 days, it is desirable to further prolong PHH functions *in vitro* towards modeling chronic drug exposure and disease progression. Fasting has benefits on the longevity of organisms and the health of tissues such as the liver. We hypothesized that a culturing protocol that mimics dynamic fasting/starvation could activate starvation pathways and prolong PHH functional lifetime. To mimic starvation, serum and hormones were intermittently removed from the culture medium of micropatterned co-cultures (MPCC) containing PHHs organized onto collagen domains and surrounded by 3T3-J2 murine fibroblasts. A weekly 2-day starvation optimally prolonged PHH functional lifetime for 6+ weeks in MPCCs versus a decline after 3 weeks in non-starved controls. The 2-day starvation also enhanced the functions of PHH monocultures for 2 weeks, suggesting direct effects on PHHs. In MPCCs, starvation activated adenosine monophosphate activated protein kinase and restricted fibroblast overgrowth onto PHH islands, thereby maintaining hepatic polarity. The effects of starvation on MPCCs were partially recapitulated by activating adenosine monophosphate activated protein kinase using metformin or growth-arresting fibroblasts via mitomycin-C. Lastly, starved MPCCs demonstrated lower false positives for drug toxicity tests and higher drug-induced cytochrome-P450 activities versus non-starved controls even after 5 weeks. In conclusion, intermittent serum/hormone starvation extends PHH functional lifetime towards enabling clinically-relevant drug screening.

---

Since drug-induced liver injury (DILI) remains a leading cause of acute liver failures (1), preclinical DILI detection is critical to mitigate the risk to patients. However, often there is a significant lack of concordance between animals and humans in DILI outcomes due to species-specific differences in drug metabolism (2–4). Furthermore, animal models of liver diseases often show significant differences in disease initiation/progression than humans, which restricts animal use for developing therapies (5, 6). In contrast, *in vitro* human liver models can mitigate the limitations with animals and enable cheaper/faster drug screening. Transformed hepatic cell lines and hepatocyte-like cells derived from induced pluripotent stem cells typically suffer from low drug metabolism capacities (7, 8), while liver slices lose viability within days (9). Therefore, cultures of primary human hepatocytes (PHHs) are ideal for drug testing; however, PHHs rapidly de-differentiate when cultured in 2-dimensional (2D) monocultures (10).

Owing to the limitations with conventional PHH monocultures, several advanced culture techniques have been developed to prolong PHH functional lifetime to 2-4 weeks *in vitro*, especially upon co-culture with liver- or non-liver-derived non-parenchymal cells (9). These include micropatterned co-cultures (MPCC), spheroids, bioprinted tissues, and microfluidic devices. The improvements in functional PHH lifetime have led to 2-3X increases in the sensitivity of DILI prediction relative to conventional monocultures and enabled the modeling of key phenotypes in liver diseases (9). However, the functional lifespan of hepatocytes ranges from 200 to 400 days *in vivo* (11, 12). Therefore, there remains a need to further prolong PHH functional lifetime *in vitro* towards modeling chronic drug exposure and disease progression.

Most PHH culture platforms utilize nutrient/hormone-rich culture medium, often containing 10% serum, for a period of 1-2 days followed by exchange with fresh medium of the same composition. The continual presence of excess nutrients/hormones in culture medium does not mimic *in* vivo fluctuations in the levels of these factors and may be partly responsible for a premature loss of PHH functions *in vitro*. Indeed, we have shown that the use of a high glucose medium (>10 mM) with a 2-day medium exchange alters drug metabolism pathways and causes insulin resistance in PHHs (13). In contrast, energy metabolism in animals is cyclical, arising from both cell autonomous circadian rhythms and the feeding-fasting cycle that modulates the activities of key regulators of nutrient homeostasis such as adenosine monophosphate activated protein kinase (AMPK) (14). The liver is especially sensitive to dynamic changes in hormones and nutrients (15) and modulating the availability of these factors in a time-restricted manner can protect rodents fed a high fat diet against obesity, hyperinsulinemia, hepatic steatosis, and inflammation (16). *In vitro,* serum starvation and hormone fluctuations have been shown to initiate fasting-stimulated signaling pathways in cell cultures and maintain normal oscillations in hepatocyte gene expression, respectively (17, 18). However, it remains unclear whether such culture modifications are useful for extending PHH functional lifetime *in vitro* and thus utility for drug development.

Here, we hypothesized that periodically starving PHH cultures of serum and hormones could maintain liver functions for a longer duration. To test our hypothesis, we utilized well-established MPCCs that maintain hepatic functions for ~3 weeks via the precise organization of PHHs onto circular collagen-coated domains of empirically optimized dimensions and co-cultivation with 3T3-J2 murine embryonic fibroblasts that express liver-like molecules to support hepatocytes (10, 19); conventional PHH monocultures were used to determine the effects on PHHs without fibroblasts. Cultures were starved of serum and hormones intermittently over 1-4 weeks and hepatic phenotype was assessed via morphological observations, albumin and urea secretions, cytochrome-P450 (CYP) enzyme activities, and the formation of functional bile canaliculi relative to non-starved control cultures. Finally, the utility of starved MPCCs for drug-induced hepatotoxicity and drug-mediated CYP induction assays was assessed.

## METHODS

### Hepatocyte culture

Cryopreserved primary human hepatocytes (PHHs) were obtained from BioIVT (Baltimore, MD) or Lonza, Inc. (Walkersville, MD): YEM (46-year-old Caucasian female, BioIVT), HUM4011 (26-year-old Caucasian male, Lonza), HUM4020 (44-year-old Caucasian male, Lonza), HUM4167 (12-year old Caucasian female, Lonza), and HUM4055A (54-year-old Caucasian female, Lonza). Micropatterned co-cultures (MPCCs) were created as previously described (20). Briefly, rat tail collagen-I (Corning, Tewksbury, MA) was lithographically patterned to create 500 μm diameter circular domains spaced 1200 μm apart, center-to-center. PHHs selectively attached to the collagen domains leaving ~25,000 attached PHHs on ~90 collagen-coated islands or ~3,500 PHHs on ~13 islands within each well of a 24-well or 96-well plate, respectively. 3T3-J2 murine embryonic fibroblasts were seeded the next day at 3:1 ratio with PHHs (**Fig. 1a**). For certain experiments, the fibroblasts were growth arrested by incubating with 1 μg/mL mitomycin-C (Sigma-Aldrich, St. Louis, MO) for 4 hours prior to co-culture. For monocultures, 350,000 PHHs were seeded in collagen-coated wells of a 24-well plate. Hepatocyte maintenance culture medium containing a Dulbecco’s Modified Eagle’s Medium (DMEM, Corning) base supplemented with 5 mM glucose, 10% bovine serum (Thermofisher, Waltham, MA), 1% penicillin/streptomycin (Corning), 1% ITS+ (insulin, transferrin, selenous acid, linoleic acid, bovine serum albumin) (Corning), 7 ng/mL glucagon (Sigma-Aldrich), 0.1 mM dexamethasone (Sigma-Aldrich), and 15 mM HEPES (Corning) was replaced on cultures every 2 days.

**Fig 1.**
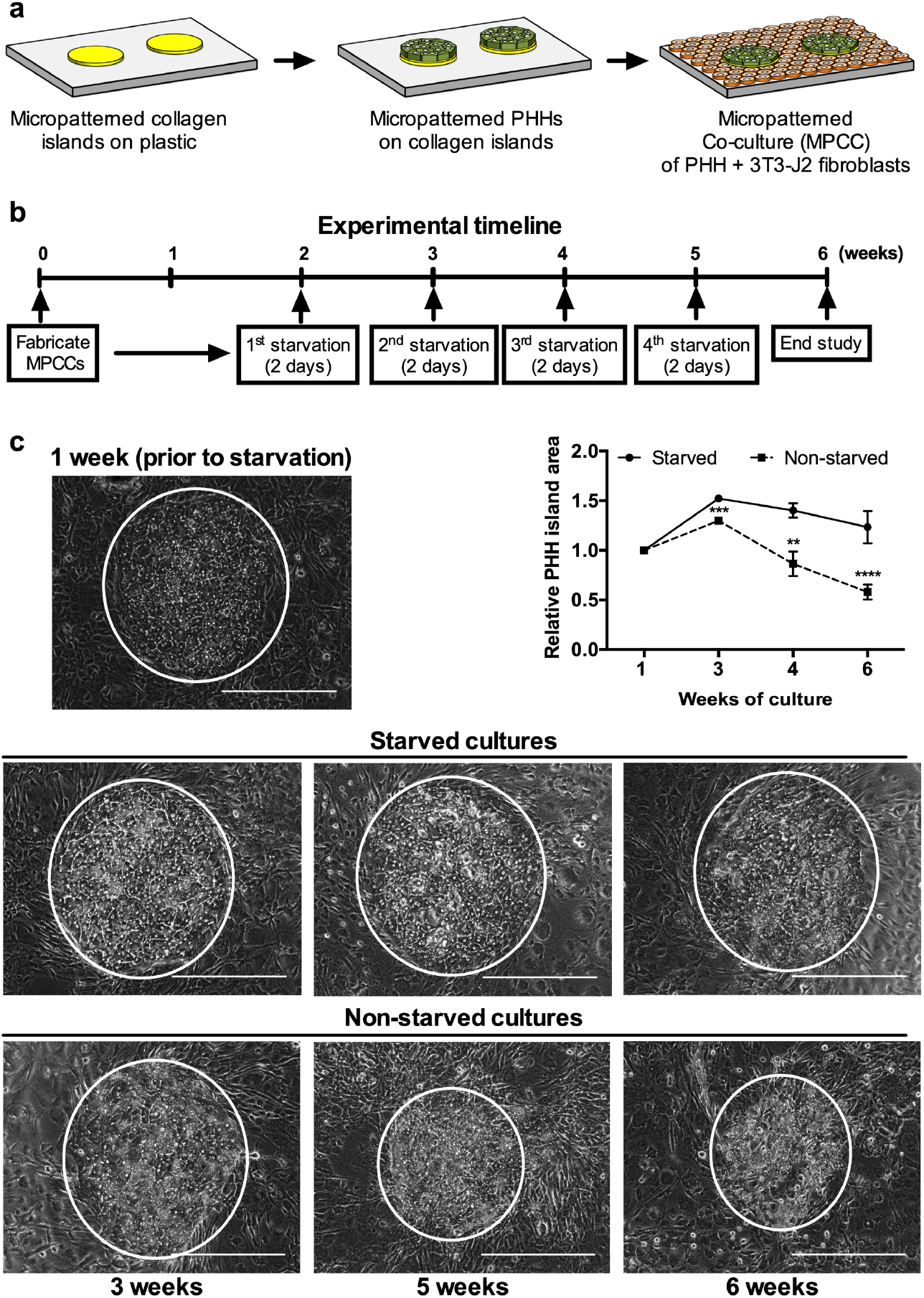
Intermittent starvation maintains hepatocyte island area and morphology over time. (a) Schematic of the MPCC platform. Left to right: Collagen-I is micropatterned using soft lithographic techniques. Hepatocytes attach to the collagen domains and unattached cells are washed off. The next day, 3T3-J2 fibroblasts are seeded in the surrounded areas. (b) Experimental timeline for intermittently starving cultures of serum and hormones (dexamethasone, glucagon, and insulin). (c) Phase contrast images of PHHs in MPCCs at 1 week prior to starvation and then at 3, 4, or 6 weeks following the weekly 2-day starvation protocol shown in panel ‘b’. The non-starved MPCC images are shown for comparison. White circles outline PHH islands. Graph on right quantifies relative PHH island area over time, normalized to PHH island area in 1-week old MPCCs. Scale bars on images represent 400 μm. \*\**p* ≤ 0.01, \*\*\**p* ≤ 0.001, and \*\*\*\**p* ≤ 0.0001.

### Intermittent starvation

MPCCs were cultured in a serum/hormone-supplemented maintenance medium for 2 weeks. Then, cultures were washed once with 1X phosphate buffered saline (Corning) and starved of all media components above except for DMEM with 5 mM glucose, 15 mM HEPES, and 1% penicillin/streptomycin for 1 hour, 1 day, 2 days, or 3 days, after which MPCCs were placed back in the serum/hormone-supplemented maintenance medium. MPCCs were subjected to starvation each week for 4 weeks. PHH monocultures were starved 2 days after seeding and every 2 days after that for 14 days. Finally, to mimic AMPK activation due to starvation, MPCCs were incubated with metformin (250 μM) in serum/hormone-containing medium every 2 days for 4 weeks.

### Biochemical assays

Albumin was measured using a competitive enzyme-linked immunosorbent assay with horseradish peroxidase detection and 3,3’,5,5’-tetramethylbenzidine (TMB, Rockland Immunochemicals, Boyertown, PA) as the substrate (10). Urea was measured using a colorimetric end-point assay utilizing diacetyl monoxime with acid and heat (Stanbio Labs, Boerne, TX) (10). CYP3A4 and CYP2C9 activities were measured by incubating cultures with luciferin-IPA or luciferin-H substrates (Promega, Madison, WI) for 1 or 3 hours, respectively. The metabolite, luciferin, was quantified via luminescence detection. CYP1A2 and CYP2A6 activities were measured by incubating cultures with 5 μM 7-ethoxyresorufin or 50 μM coumarin (Sigma-Aldrich) for 1 hour, respectively. The metabolites, resorufin and 7-hydroxycoumarin (7-HC), generated from 7-ethoxyresorufin and coumarin, respectively, were quantified via fluorescence detection (excitation/emission: 550/585 nm for resorufin, and 355/460 nm for 7-HC) (20).

### Cell imaging

Culture morphology was monitored using an EVOS^®^ FL microscope (Thermofisher). Additionally, cultures were first washed 1X with serum-free culture medium and then incubated for 15 minutes with 2 μM of 5(6)-carboxy-2’,7’-dichlorofluorescein-diacetate (CDCFDA) that gets cleaved into 5(6)-carboxy-2’,7’-dichlorofluorescein (CDF) by esterases in hepatocytes and exported into the bile canaliculi by multidrug resistance protein-2 (21) and 1 μM of Hoechst-33342 that stains DNA. Cultures were then washed 3X with a serum-free culture medium and imaged using the GFP (green fluorescent protein) and DAPI (4’,6-diamidino-2-phenylindole) light cubes to visualize the CDF and nuclear stains, respectively. Analysis of hepatocyte island area and solidity was carried out in Fiji (22) by outlining phase contrast images of hepatocyte islands and using the “shape descriptors” feature in the analysis. The area of excreted CDF was assessed by setting a threshold that included all fluorescent signal and subsequently quantifying the threshold area per PHH island.

### Drug studies

Drugs were purchased from Sigma-Aldrich or Cayman Chemicals (Ann Arbor, MI) and dissolved in 100% dimethylsulfoxide (DMSO, Corning). MPCCs were cultured for 4 weeks +/− intermittent starvation as above. Then, the cultures were treated with hepatotoxic (diclofenac, troglitazone, piroxicam, amiodarone, clozapine) or non-hepatotoxic drugs (rosiglitazone, prednisone, miconazole, aspirin, dexamethasone) dissolved in serum-free culture medium at 25× and 100× C_max_ (C_max_: maximum drug concentration measured in human plasma) for each drug; cultures were treated with drugs for 6 days with fresh drug added to culture medium at each 2-day medium exchange.

Drug/C_max_ (μM): amiodarone/0.806, aspirin/5.526, clozapine/0.951, dexamethasone/0.224, diclofenac/8.023, miconazole/0.024, piroxicam/5.135, prednisone/0.068, rosiglitazone/1.120, troglitazone/6.387 (23). Following drug exposure, albumin and urea secretions, previously shown to correlate highly with hepatotoxicity markers such as ATP (3), were measured as above. DMSO concentration in the medium was kept at 0.1% (v/v), and a DMSO-only control culture was used to calculate relative changes in endpoints due to drug treatment.

For CYP induction, intermittently starved MPCCs and non-starved controls were exposed to serum-free culture medium containing rifampin (25 μM), phenobarbital (1 mM), omeprazole (10 μM), or dimethylsulfoxide (DMSO) alone (0.1% v/v); cultures were treated with drugs for 4 days with fresh drug added to culture medium at the 2-day medium exchange. Following drug exposure, CYP3A4, CYP2C9, and CYP1A2 activities were quantified as above.

### Data analysis

All findings were confirmed in 3-4 wells per condition across different PHH donors. Data processing was performed using Microsoft Excel. GraphPad Prism (La Jolla, CA) was used for displaying results. Mean and standard deviation are displayed for all data sets for each data point. Statistical significance was determined using Student’s t-test (two groups) or one-way ANOVA followed by a Dunnett’s multiple comparison test (p< 0.05) for more than two groups. For drug toxicity studies, IC_50_ values (drug concentrations that reduced an end-point by 50% of DMSO-only controls) were calculated by the linear interpolation between the drug dose at which the assay signal was >50% of controls and the drug dose at which the assay signal was <50% of controls.

## RESULTS

### Optimization of the intermittent starvation protocol

MPCCs were cultured in serum/hormone-supplemented medium for 2 weeks and then subjected to serum/hormone-free medium for 1 hour, 1 day, 2 days, or 3 days, after which MPCCs were placed back in the serum/hormone-supplemented medium; this intermittent starvation protocol was repeated on the cultures each week for 4 weeks. Albumin and urea secretions were measured in supernatants while the presence/functionality of the bile canaliculi was assessed using export of the CDF dye through hepatic transporters. Our results showed that the PHHs better retained prototypical morphology with the 2-day and 3-day intermittent starvations as compared to the 1-hour and 1-day starvations and the non-starved controls (**Supplemental Fig. 1a**). At the functional level, all starvation durations led to higher albumin than non-starved controls, though the 1-hour starvation had the lowest urea while the 1-day, 2-day and 3-day starvations had statistically similar urea (**Supplemental Fig. 1b**). In contrast, the 1-day starvation led to the highest urea as compared to other starvation durations and the non-starved controls (**Supplemental Fig. 1c**). Lastly, the functional bile canaliculi network was enhanced with the 1-day, 2-day, and 3-day starvations as compared to the 1-hour starvation and non-starved controls, with 2-day starvation being the optimal (**Supplemental Fig. 1d**). Therefore, since two of the three functional markers (albumin and bile canaliculi) were retained at the highest level and morphology was maintained over time, we selected the 2-day duration for intermittent starvations of cultures for the remainder of our studies.

### Intermittent starvation leads to long-term retention of PHH morphology and function

MPCCs that were 2 weeks old were subjected to 2-day starvations for 4 additional weeks (**Fig. 1b**). The overall area of PHH islands and prototypical hepatic morphology were retained for 6 weeks in intermittently starved cultures whereas these features degraded in non-starved cultures (**Fig. 1c**). At the functional level, intermittently starved MPCCs outperformed non-starved controls over several weeks in culture and the functional differences between the two conditions increased over time for several endpoints. Specifically, over 6 weeks, albumin in the intermittently starved cultures was 1.6- to 3.6-fold higher (**Fig. 2a**), urea was 1.5- to 7.7-fold higher (**Fig. 2b**), CYP3A4 activity was 1.4- to 14.3-fold higher (**Fig. 2c**), and CYP2A6 activity was ~2-fold higher (**Supplemental Fig. 2**) than non-starved controls.

**Fig 2.**
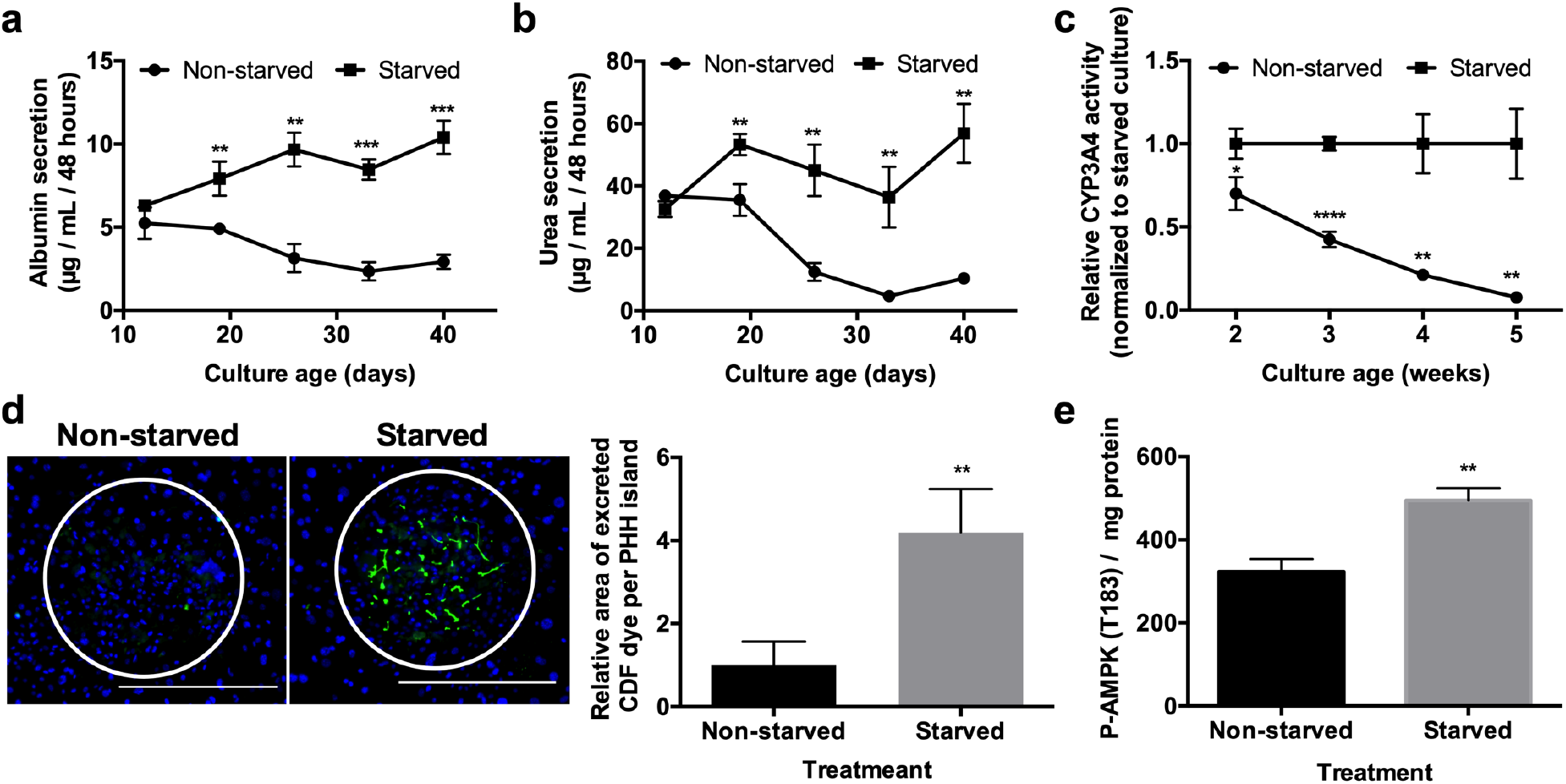
Intermittent starvation maintains hepatocyte function and causes AMPK activation. (a) Albumin and (b) urea secretions, and (c) CYP3A4 enzyme activity in starved (as described in Figure 1b) and non-starved cultures over time. (d) Functional bile canaliculi in PHHs within starved and non-starved MPCCs as assessed by the excretion of the CDF dye. Left: images of representative bile canaliculi in PHH islands within the two conditions. Right: Relative area of excreted CDF in PHH islands within starved versus non-starved MPCCs after 6 weeks of culture. Data is normalized to the non-starved controls. (e) Phosphorylated-AMPK (p-AMPK) per mg of protein in MPCC lysates following 2 starvation periods. Scale bars on images represent 400 μm. *p<0.05, \*\**p* ≤ 0.01, \*\*\**p* ≤ 0.001, and \*\*\*\**p* ≤ 0.0001.

Additionally, after 6 weeks, the formation of functional bile canaliculi in PHHs within MPCCs was ~4-fold higher in intermittently starved cultures than non-starved controls (**Fig. 2d**). Such an outcome correlated with an increase of total phosphorylated adenosine monophosphate activated kinase (p-AMPK) in intermittently starved MPCCs relative to the non-starved controls (**Fig. 2e**); p-AMPK has been previously implicated in maintaining hepatocyte polarity (24, 25).

We also tested the intermittent starvation protocol on conventional 2D PHH monocultures but only over 14 days owing to their shorter functional lifetime; PHH monocultures were starved 2 days after seeding and every 2 days thereafter. As in MPCCs, PHH morphology and confluency were better retained in intermittently starved PHH monocultures than non-starved controls (**Supplemental Fig. 3a**). Furthermore, intermittently starved PHH monocultures showed improvement of several functions over non-starved controls. Specifically, intermittently starved cultures displayed higher CYP activities (1.7 fold for CYP2A6 and 2.7 fold for CYP3A4, **Supplemental Fig. 3b**) and a greater number of functional bile canaliculi (**Supplemental Fig. 3c**) after 2 weeks than non-starved controls. However, in contrast to MPCCs, albumin and urea were only marginally (not statistically significant) improved in intermittently starved PHH monocultures than non-starved controls (**Supplemental Fig. 3).**

**Fig 3.**
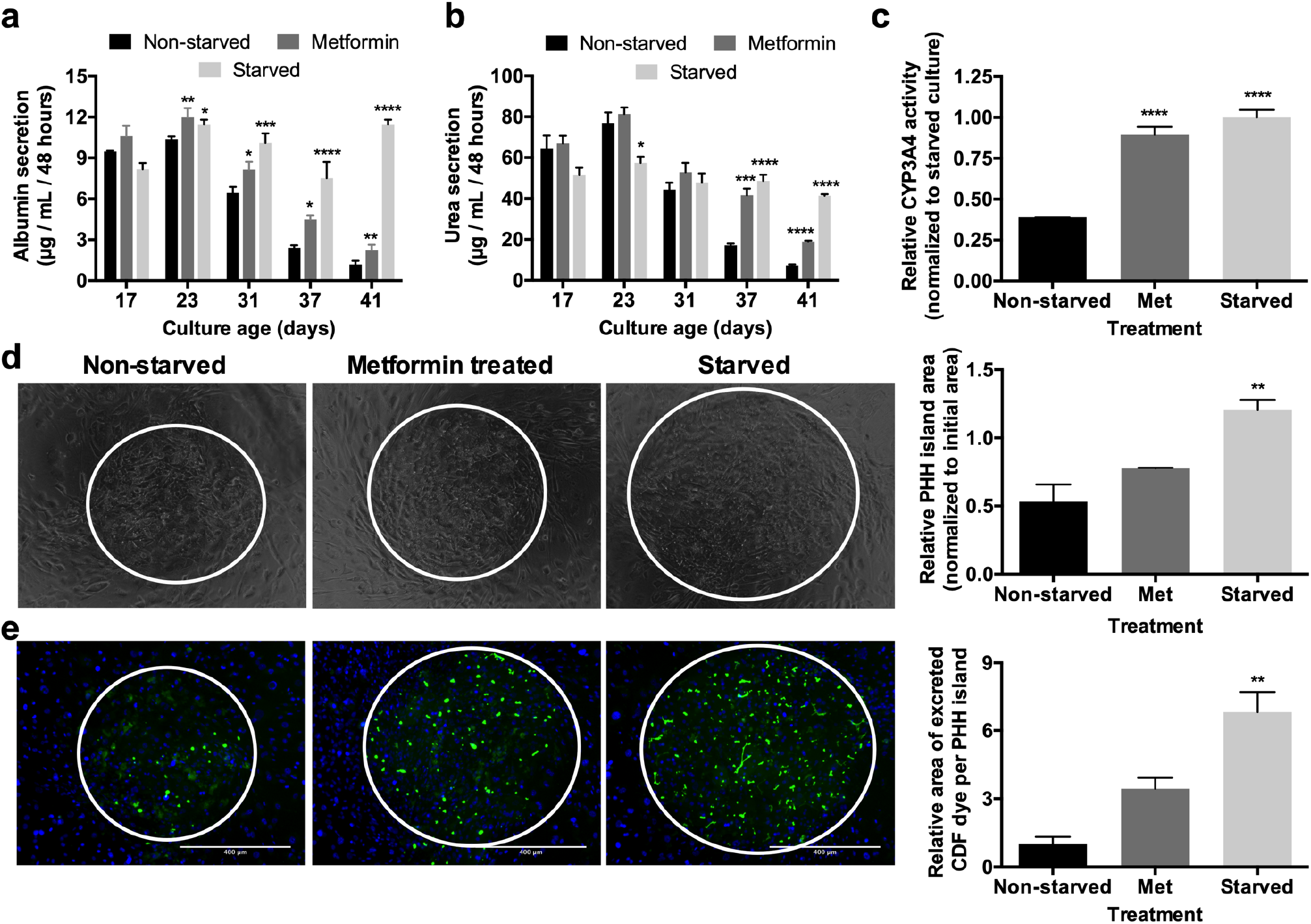
Metformin treatment recapitulates some of the functional benefits of intermittent starvation. (a) Albumin production over time, (b) urea synthesis over time, and (c) CYP3A4 enzyme activity after 6 weeks of culture in non-starved, starved (as described in Figure 1b), and metformin treated MPCCs. Metformin treatment followed the same schedule as starvation except that cultures were treated with metformin in serum/hormone-supplemented culture medium instead. (d) Phase contrast images of MPCCs under various treatments as described above after 6 weeks of culture. White circles outline PHH islands. Graph on the right quantifies relative PHH island area in MPCCs under various treatments after 6 weeks of culture. (e) Functional bile canaliculi in PHHs within MPCCs under various treatments as assessed by the excretion of the CDF dye. Left: images of representative bile canaliculi in PHH islands within the three conditions. Right: Relative area of excreted CDF in PHH islands within the 3 conditions after 6 weeks of culture. Data is normalized to the non-starved controls. Scale bars on images represent 400 μm. *p<0.05, \*\**p* ≤ 0.01, \*\*\**p* ≤ 0.001, and \*\*\*\**p* ≤ 0.0001 relative to the non-starved control.

### Metformin treatment recapitulates some of the functional benefits of intermittent starvation

Since we observed an increase in p-AMPK in intermittently starved MPCCs (**Fig. 2e**), we assessed the effects of pharmacologic AMPK activation via metformin in lieu of intermittent starvation. We treated MPCCs with metformin for 2 days every week for 4 weeks in serum/hormone-containing medium. Metformin-treated MPCCs displayed ~1.9-fold higher albumin (**Fig. 3a**), ~2.6 higher urea (**Fig. 3b**), and ~2.3-fold higher CYP3A4 activity (**Fig. 3c**) after ~6 weeks of culture as compared to non-starved controls. However, such functional upregulation in metformin-treated MPCCs was still lower than in intermittently starved MPCCs (~9.7-fold, ~5.7-fold, ~2.6-fold higher in starved vs. non-starved cultures after 6 weeks for albumin, urea, and CYP3A4, respectively). Similarly, metformin-treated MPCCs displayed a marginally higher (~1.5-fold), albeit statistically insignificant, PHH island area relative to the non-starved controls, which was lower than the ~2.3-fold higher PHH island area achieved in the starved MPCCs (**Fig. 3d**). Finally, metformin-treated MPCCs displayed an enhanced network of functional bile canaliculi (~3.4-fold higher area of excreted CDF dye) relative to the non-starved controls, which was lower than the ~6.8-fold higher area of excreted CDF dye achieved in the starved MPCCs (**Fig. 3e**).

### Intermittent starvation prevents fibroblast overgrowth

Besides the PHH specific changes with intermittent starvation, we noticed changes in fibroblast morphology during starvation and subsequent cell density following starvation. During starvation, some fibroblasts assumed a rounded, rather than spread, morphology, which suggested that select fibroblasts might detach or undergo apoptosis during the serum starvation. Using fibroblast monocultures, we found that fibroblast DNA was reduced by ~4 fold after a 2-day serum starvation period (**Supplemental Fig. 4**). We confirmed this finding in MPCCs by counting fibroblast nuclei in between PHH islands and observed a lower fibroblast nuclei count in the starved cultures relative to the non-starved controls (**Fig. 4a**). In contrast, fibroblast numbers did not change significantly in metformin-treated MPCCs relative to the non-starved controls (**Supplemental Fig. 5**).

**Fig 4.**
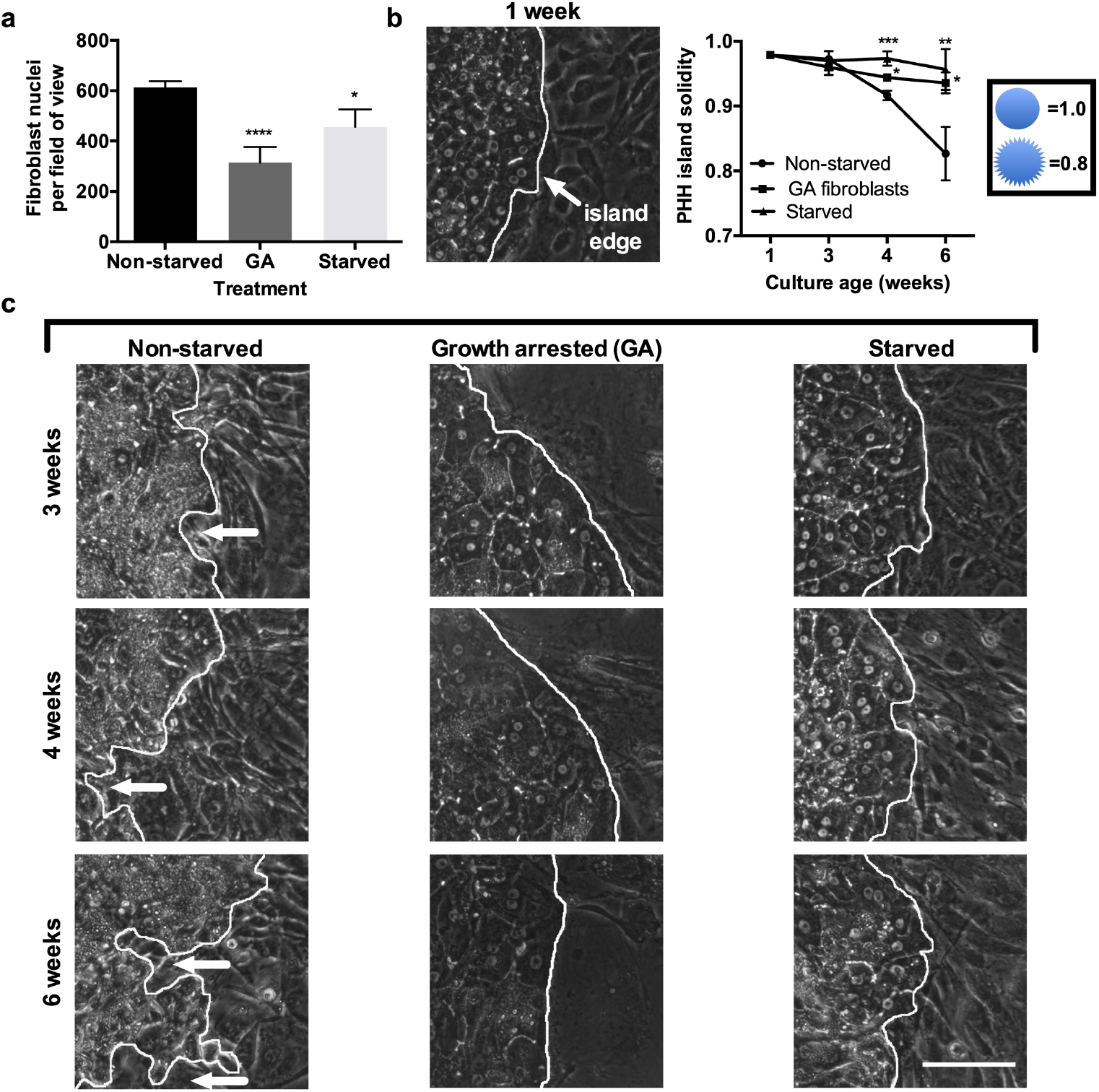
Growth arresting fibroblasts maintains hepatocyte island integrity in a similar way as intermittent starvation. MPCCs were created using either proliferative fibroblasts for the non-starved and starved (as described in Figure 1b) conditions or growth-arrested (GA) fibroblasts via mitomycin-C. (a) Fibroblast nuclei count between PHH islands in 6-week old MPCCs (3 different models described above) as assessed via live cell imaging of nuclei using Hoechst 33342 staining. (b) Phase contrast images of PHH island edges outlined in white in the three different models above over time. White arrows indicate fibroblasts invading the PHH island. The 1 week time-point shown is for the non-starved MPCCs. Graph on top right shows PHH island solidity (island area/convex hull area) in the different models over time. Scale bars on images represent 50 μm. *p<0.05, \*\**p* ≤ 0.01, \*\*\**p* ≤ 0.001, and \*\*\*\**p* ≤ 0.0001 relative to the non-starved control.

**Fig 5.**
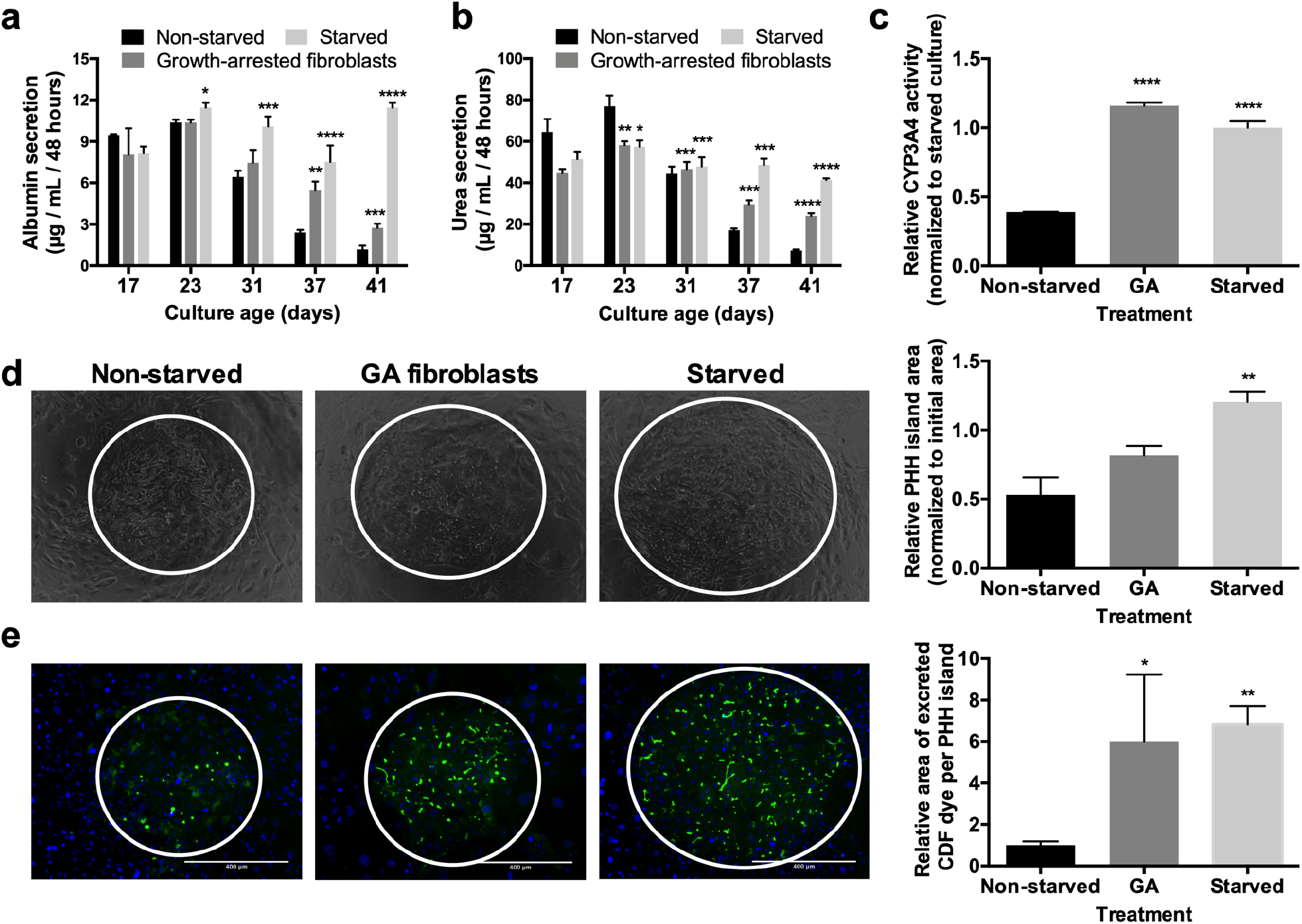
Growth arresting fibroblasts recapitulates some of the functional benefits of intermittent starvation. MPCCs were created using either proliferative fibroblasts for the non-starved and starved (as described in Figure 1b) conditions or growth-arrested (GA) fibroblasts. (a) Albumin production over time, (b) urea synthesis over time, and (c) CYP3A4 enzyme activity after 6 weeks of culture in non-starved MPCCs, starved MPCCs, and MPCCs with growth-arrested fibroblasts. (d) Phase contrast images of different MPCC models as described above after 6 weeks of culture. White circles outline PHH islands. Graph on the right quantifies relative PHH island area in different MPCC models after 6 weeks of culture. (e) Functional bile canaliculi in PHHs within different MPCC models after 6 weeks of culture as assessed by the excretion of the CDF dye. Left: images of representative bile canaliculi in PHH islands within the three MPCC models. Right: Relative area of excreted CDF in PHH islands within the different MPCC models. Data is normalized to the non-starved controls. Scale bars represent 400 μm. *p<0.05, \*\**p* ≤ 0.01, \*\*\**p* ≤ 0.001, and \*\*\*\**p* ≤ 0.0001 relative to the non-starved control. Note: The data and images for starved and non-starved conditions shown in this figure are the same as those shown in figure 3 for comparison with the GA condition.

### Growth arresting fibroblasts recapitulates some of the functional and morphological benefits of intermittent starvation

Since the findings above indicated that a reduction in fibroblast number over time may partly explain improvements in PHH functions in intermittently starved MPCCs, we growth arrested fibroblasts using mitomycin-C prior to incorporation into MPCCs to prevent fibroblast overgrowth in lieu of intermittent starvation. Growth arresting successfully reduced the number of fibroblasts in MPCCs over time as compared to the MPCCs with proliferative (non-starved) fibroblasts (**Fig. 4a**). Furthermore, the solidity of PHH islands, as assessed by dividing the PHH island area by the convex hull area, was significantly higher in MPCCs containing growth-arrested fibroblasts (non-starved) and intermittently starved MPCCs containing proliferative fibroblasts as compared to the non-starved MPCCs containing proliferative fibroblasts (**Fig. 4b**).

At the functional level, MPCCs containing growth-arrested fibroblasts displayed ~2.3-fold higher albumin (**Fig. 5a**), ~3.3 higher urea (**Fig. 5b**), and ~3-fold higher CYP3A4 activity (**Fig. 5c**) after 6 weeks of culture as compared to non-starved controls. However, upregulation of albumin and urea in MPCCs containing growth-arrested fibroblasts was still lower than observed with intermittently starved MPCCs containing proliferative fibroblasts (~9.7-fold and ~5.7-fold in starved vs. non-starved cultures after 6 weeks of culture for albumin and urea, respectively). On the other hand, CYP3A4 activities in MPCCs containing growth-arrested fibroblasts and intermittently starved MPCCs with proliferative fibroblasts were similar. Furthermore, MPCCs containing growth-arrested fibroblasts displayed a marginally higher (~1.5-fold), albeit statistically insignificant, PHH island area relative to the non-starved controls, which was lower than the ~2.3-fold higher PHH island area achieved in the starved MPCCs with proliferative fibroblasts (**Fig. 5d**). Finally, MPCCs containing growth-arrested fibroblasts displayed an enhanced network of functional bile canaliculi (~6-fold higher area of excreted CDF dye), which was slightly lower than the ~6.8-fold higher area of excreted CDF dye achieved in the starved MPCCs with proliferative fibroblasts as compared to the non-starved controls (**Fig. 5e**).

### Intermittent starvation leads to lower false positives for drug toxicity assays

In order to assess whether intermittently starved cultures can be used for drug toxicity screening, MPCCs were first cultured for up to 4 weeks +/− intermittent starvation as above. Then, the cultures were treated for a total of 6 days with 5 hepatotoxic drugs (amiodarone, clozapine, diclofenac, piroxicam, troglitazone) or 5 non-hepatotoxic drugs (aspirin, dexamethasone, miconazole, prednisone, rosiglitazone) at 25× and 100× C_max_ (C_max_: maximum drug concentration measured in human plasma (23)) for each drug, with fresh drug added at each 2-day culture medium exchange (**Fig. 6a**). Previously, 5-9 days of treatment with drugs at concentrations up to 100× C_max_ was found to increase the sensitivity for drug toxicity detection without an increase in the false positive rate over 1-day drug treatment in MPCCs (3). The DMSO concentration in the culture medium was kept at 0.1% (v/v) for all drugs since DMSO at high levels can inhibit CYP3A4 activity (26). A DMSO-only control culture was used to calculate relative changes in functional endpoints due to the drug treatment. Albumin and urea, specific to PHHs, were measured after the 6-day drug exposure period since downregulation of these functions was shown to correlate with toxicity endpoints such as ATP in PHHs (3, 27, 28). The IC_50_ values (drug concentration that reduces function by 50%) after 6 days of drug treatment were interpolated from the dose response curves for both albumin (**Supplemental Fig. 6**) and urea (**Supplemental Fig. 7**).

**Figure 6.**
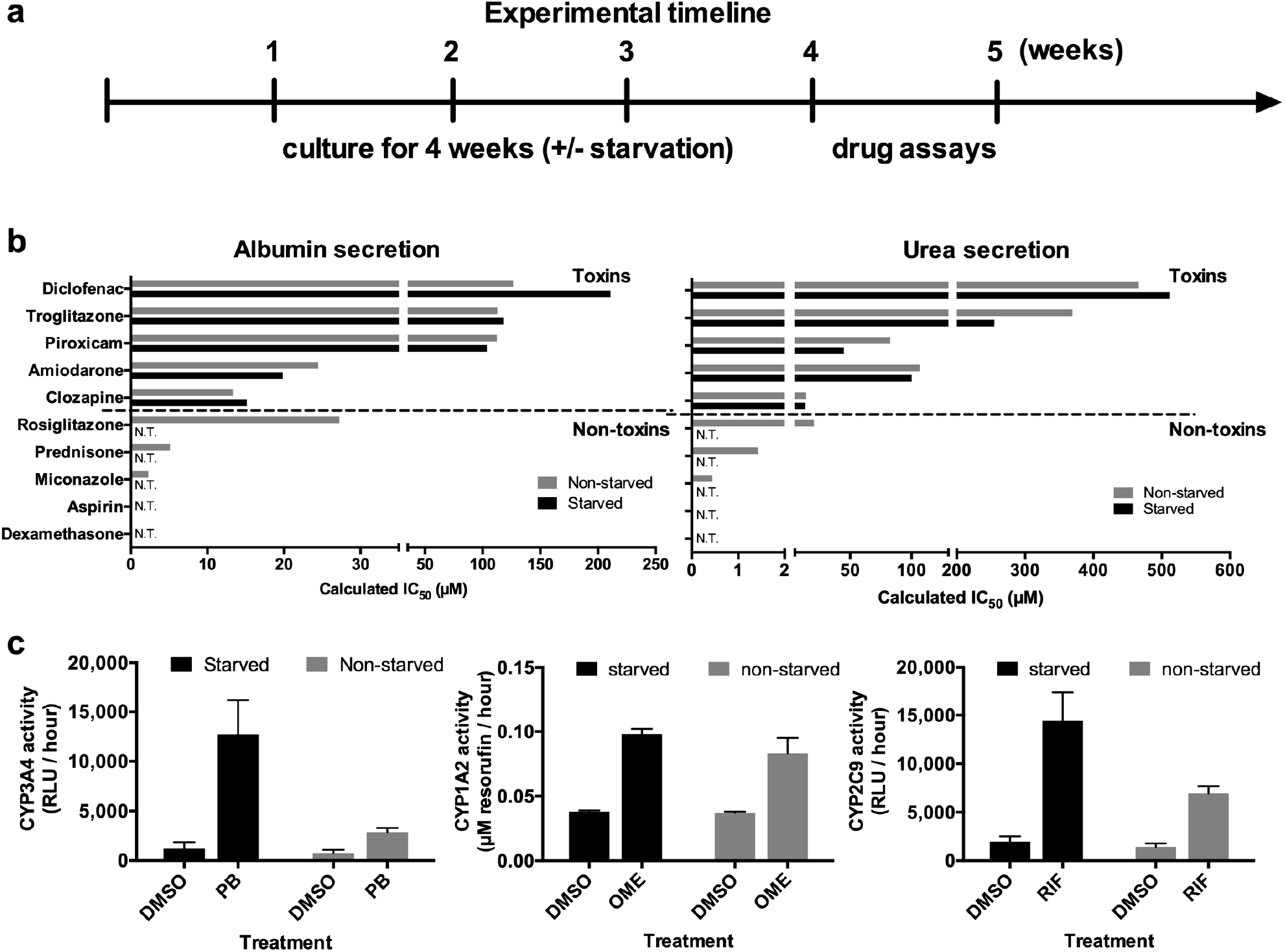
Intermittent starvation prolongs MPCC utility for clinically-relevant drug screening. (a) MPCCs were maintained either non-starved or starved for 2 days every week for 4 weeks (as described in Figure 1b) and then treated with drugs. (b) IC_50_ values for albumin (left) and urea (right) for starved and non-starved cultures treated for 6 days with prototypical hepatotoxins (top) and non-hepatotoxins (bottom) at 25× and 100× C_max_ (C_max_: maximum drug concentration measured in human plasma) for each drug. N.T. represents ‘not toxic’ (i.e. IC_50_ value could not be interpolated). (c) Induction of CYP enzymes in starved and non-starved MPCCs after 4 days of treatment with prototypical inducer drugs (PB: phenobarbital, OME: omeprazole, and RIF: rifampin).

Albumin IC_50_ values after 6 days of drug treatment were similar in starved vs. non-starved MPCCs for 4 out of the 5 hepatotoxic drugs (amiodarone, clozapine, piroxicam, troglitazone), while the albumin IC_50_ value for diclofenac was ~1.7-fold higher in starved vs. non-starved MPCCs (**Fig. 6b**). Urea IC_50_ values were similar in starved vs. non-starved MPCCs for 4 out of the 5 hepatotoxic drugs (clozapine, diclofenac, piroxicam, troglitazone), while the urea IC_50_ value for amiodarone was ~1.8-fold lower in starved vs. non-starved MPCCs (**Fig. 6b**). In spite of the differences in IC_50_ values, both starved and non-starved MPCCs correctly detected the liver dysfunctions/hepatotoxicity caused by the hepatotoxic drugs, which is consistent with previously published data 1-week old and non-starved MPCCs treated with the same compounds (3).

Neither albumin nor urea IC_50_ values could be interpolated in starved MPCCs for any of the 5 non-hepatotoxins even after 6 days of treatment, suggesting a high level of specificity of the assay (i.e. no false positives) even when cultures were ~5 weeks old (**Fig. 6b**). In contrast, albumin and urea IC_50_ values could indeed be interpolated in non-starved MPCCs after 6 days of treatment for 3 out of the 5 non-hepatotoxins (miconazole, prednisone, rosiglitazone) (**Fig. 6b**). Furthermore, rosiglitazone caused higher hepatic dysfunctions (i.e. lower IC_50_ values) than troglitazone in 5-week old non-starved MPCCs, which does not correlate with the relative clinical hepatoxicities of these compounds. Thus, the 5-week old non-starved MPCCs detected 3 out of 5 non-hepatotoxins as false positives whereas 5-week old intermittently starved MPCCs detected no false positive drugs; the high specificity of the starved MPCCs is similar to that obtained in 1-week old and non-starved MPCCs previously treated with the same drugs (3).

### Intermittent starvation leads to higher levels of drug-mediated CYP induction

In order to assess whether intermittently starved cultures can be used for drug-mediated CYP induction assessment (a measure of drug-drug interactions), MPCCs were first cultured for up to 4 weeks +/− intermittent starvation as above. Then, the cultures were treated for a total of 4 days with single concentration of either a CYP1A2 inducer (omeprazole), CYP3A4 inducer (phenobarbital), or a CYP2C9 inducer (rifampin), with fresh drug added every 2 days. Control cultures were treated with DMSO alone. The activities of the corresponding CYP enzymes were assessed as detailed in methods.

The starved MPCCs displayed higher CYP activities for both baseline (DMSO treated) and drug-treated conditions as compared to non-starved controls (**Fig. 6c**). The fold increases in CYP activities in drug-treated conditions relative to the DMSO-treated controls were also greater in starved MPCCs. Phenobarbital-mediated CYP3A4 induction was 10.2 fold and 3.7 fold in starved and non-starved MPCCs, respectively. Rifampin-mediated CYP2C9 induction was 7.3 fold and 4.8 fold in starved and non-starved MPCCs, respectively. Finally, omeprazole-mediated CYP1A2 induction was 2.6 fold and 2.2 fold in starved and non-starved MPCCs, respectively.

## DISCUSSION

Recent advances in engineered culture platforms can prolong the lifetime of PHHs *in vitro*. Successful efforts have taken inspiration from *in vivo* microenvironmental cues (9); however, the impact of dynamic nutrient/hormonal fluctuations on hepatocyte longevity has not been investigated. Here, we sought to address the potential benefits of mimicking the fasting/feeding cycles in the body on hepatocyte functional lifetime *in vitro*. We developed a protocol whereby serum and hormones are periodically removed from the culture medium to mimic key aspects of fasting/feeding in the liver, which we termed intermittent starvation (29). We found that intermittent starvation every week had profound effects on hepatocytes ranging from morphologic stability, prolonged hepatocyte function, and clinically-relevant responses to drugs.

Since hepatocytes rapidly de-differentiate in 2D monocultures, we utilized the MPCC platform which has been shown to sustain hepatocyte functions from multiple species for 3-4 weeks via supportive factors from 3T3-J2 murine embryonic fibroblasts; the fibroblasts are essential as none of the liver-derived non-parenchymal cells (i.e. liver sinusoidal endothelial cells, hepatic stellate cells, and Kupffer cells) support hepatocytes to the same degree as the fibroblasts (9). Importantly, PHHs in MPCCs appropriately respond to hormones and nutrients, which is necessary to investigate the potential utility of dynamic culturing methods such as intermittent starvation (13). Furthermore, precisely patterned PHH colonies can be tracked over time in MPCCs to assess the maintenance of PHH homotypic contact and formation of functional bile canaliculi. Nonetheless, to determine the generalizability of our findings and to elucidate effects on PHHs alone, we also tested intermittent starvation on PHH monocultures.

The MPCC platform necessitates a maintenance medium comprised of 5 mM glucose, 10% bovine serum, and hormones (insulin, glucagon, dexamethasone); the nutrients and growth factors in serum are essential for proper fibroblast morphology/growth, which in turn affects PHH phenotype. Typically, the culture medium is exchanged every 1-2 days in MPCCs, which is similar to other PHH culture platforms. However, during fasting *in vivo*, nutrients and insulin levels are known to decrease (30). We therefore mimicked aspects of such fasting by intermittently removing serum and hormones from our culture medium. After 2 weeks of stabilization in serum/hormone-supplemented medium, we incubated MPCCs with a culture medium lacking serum or hormones for variable time periods (1 hour, 1 day, 2 days, 3 days), after which the MPCCs were placed back in the serum/hormone-supplemented medium; this starvation protocol was repeated each week for 4 weeks. Glucose (5 mM) was kept in the starvation medium since glucose levels are highly regulated in blood; furthermore, removal of glucose from the starvation medium led to a premature decline of PHH function (not shown).

Surprisingly, we found that intermittent starvation increased the functional lifetime of PHHs in MPCCs. While all starvation durations, including 1-hour, had some benefits on hepatocyte morphology and functions, the 2-day starvation period was optimal. MPCCs that were intermittently starved for 2-days every week for 4 weeks had similar morphology and island size as 2-week old cultures, whereas PHH islands shrunk in size in the non-starved controls. Additionally, albumin and urea in starved MPCCs were sustained at similar or higher levels as 2-week old cultures whereas function of non-starved MPCCs rapidly declined after 3 weeks. Additionally, CYP activity was retained to at least 50% of the 2-week values in starved cultures after 5 weeks, while non-starved cultures lost most of their CYP activity by that time. Furthermore, the retention of functional bile canaliculi was enhanced in starved cultures versus non-starved cultures after 4 weeks of intermittent starvation. PHH monocultures also showed functional improvements when subjected to 2-day starvations every two days for 14 days. Therefore, intermittent starvation is useful to enhance PHH functional lifetime in different model systems.

Periodic fasting is associated with increased lifetime in yeast, rodents, and humans, and has been attributed to the activation of the energy sensing enzyme AMPK (31–33). Serum starvation is known to activate this protein in cell cultures (18); we also found that our starvation protocol led to increased levels of activated (phosphorylated) AMPK. Metformin can also activate AMPK (24, 34); thus, we used periodic metformin treatment (in lieu of starvation) to identify the contribution of AMPK activation on prolonging hepatocyte functions. While metformin treatment prolonged some PHH functions, it did not improve either the retention of PHH island size or the functional bile canaliculi network. Thus, AMPK activation is necessary but not sufficient to yield the entire spectrum of functional benefits observed with intermittent starvation.

It has previously been reported that supportive fibroblasts in co-culture with epithelial cells must be prevented from overgrowing the epithelial population to prevent a decline in epithelial cell colony integrity (35). In intermittently starved MPCCs, we observed a reduction in fibroblast numbers in between PHH islands, which also led to less encroachment of the fibroblasts onto the PHH islands over several weeks and thereby better retained PHH homotypic contacts. To test whether a reduction in fibroblast number had an effect on better retention of PHH functions, we growth arrested fibroblasts via mitomycin-C prior to co-culture with PHHs. Growth-arresting the fibroblasts improved the functional output of non-starved MPCCs as compared to non-starved MPCCs containing proliferative fibroblasts over 6 weeks. However, overall, such improvements were still at a lower level than starved MPCCs containing proliferative fibroblasts. These results coupled with the metformin results above suggest that the effects of starvation are likely multifactorial, involving both direct effects on PHHs, fibroblasts, and reciprocal hepatocyte-fibroblast interactions.

In addition to AMPK activation and restrictions in fibroblast numbers, intermittent starvation may have other effects on hepatocyte longevity. Abnormal hormone levels and an abundance of nutrients, which are present in standard culture medium, are associated with altered liver function, the buildup of toxins and metabolic dysfunctions (15–17, 29). Therefore, better retention of CYPs and transporters in the intermittently starved MPCCs may enable enhanced removal of built-up toxins over time in the PHHs. Lastly, we observed that PHHs accumulated fewer lipid droplets when exposed to the intermittent starvation protocol, which could limit the buildup of toxic lipids and prevent premature cell apoptosis (36).

Long-term PHH cultures are useful to assess the potential for drug-induced hepatotoxicity and drug-drug interactions in preclinical pharmaceutical development. The MPCC platform (non-starved) has been validated for use in such applications with sensitivities that are 2-3X higher than in conventional PHH monocultures. However, drug treatments are typically initiated after 1-week of culture when the PHHs are highly functional in MPCCs and then carried out over ~1 additional week when the cultures are 2 weeks of age. Whether such drug treatments can be carried out at later culture ages has yet to be determined; such a capability is highly desirable for pharmaceutical practice to a) model chronic drug effects and b) extend culture shelf-life towards staggering testing of a larger compound set. Therefore, we tested the utility of intermittently starved MPCCs for drug-induced hepatotoxicity and drug-mediated CYP induction assays after 4 weeks of culture and compared to non-starved controls of the same culture age. Of the 5 hepatotoxic drugs, both the non-starved and starved MPCCs correctly classified these compounds as toxic based on downregulations of albumin and urea secretions, which correlate with compound-induced hepatotoxicity (as assessed with markers such as ATP) in MPCCs and other platforms (3, 28, 37). However, for the 5 non-hepatotoxic drugs, non-starved MPCCs falsely identified 3 of 5 drugs as toxic, whereas starved MPCCs correctly identified all 5 compounds as non-hepatotoxic, which is consistent with previously published results with treatment of 1-week old and non-starved MPCCs with the same compound set (3).

For drug-mediated CYP induction, intermittently starved MPCCs displayed higher fold increases in CYP activities in drug-treated conditions relative to DMSO-treated controls as compared to non-starved MPCCs; such increases are desirable for pharmaceutical practice towards providing a larger dynamic range for rank ordering compounds based on their variable propensity for inducing CYP activities. Therefore, our drug studies suggest that intermittently starved MPCCs respond to drugs in a clinically-relevant manner even after 5 weeks of culture.

In conclusion, we developed an intermittent starvation protocol to extend the functional lifetime of PHHs towards enabling highly sensitive and specific assessment of drug responses even after 5 weeks of culture. Lastly, we showed that AMPK activation and restriction in fibroblast growth/numbers partly underlie the observed effects of intermittent starvation. We anticipate that our protocol can find broad utility in the continued development of human liver models for drug development and ultimately, cell-based therapies.

## Acknowledgements

Funding was provided by the National Institutes of Health (1R03EB019184-01 to SRK) and the National Science Foundation (CBET-1351909 to SRK). The authors would like to thank Christine Lin, Adam Lejune, Josh Pickrell, Jaron Thompson, and Brenton Ware for cell culture assistance.

