## Supplemental Data for "Intermittent Starvation Extends the Functional Lifetime of Primary Human Hepatocyte Cultures"

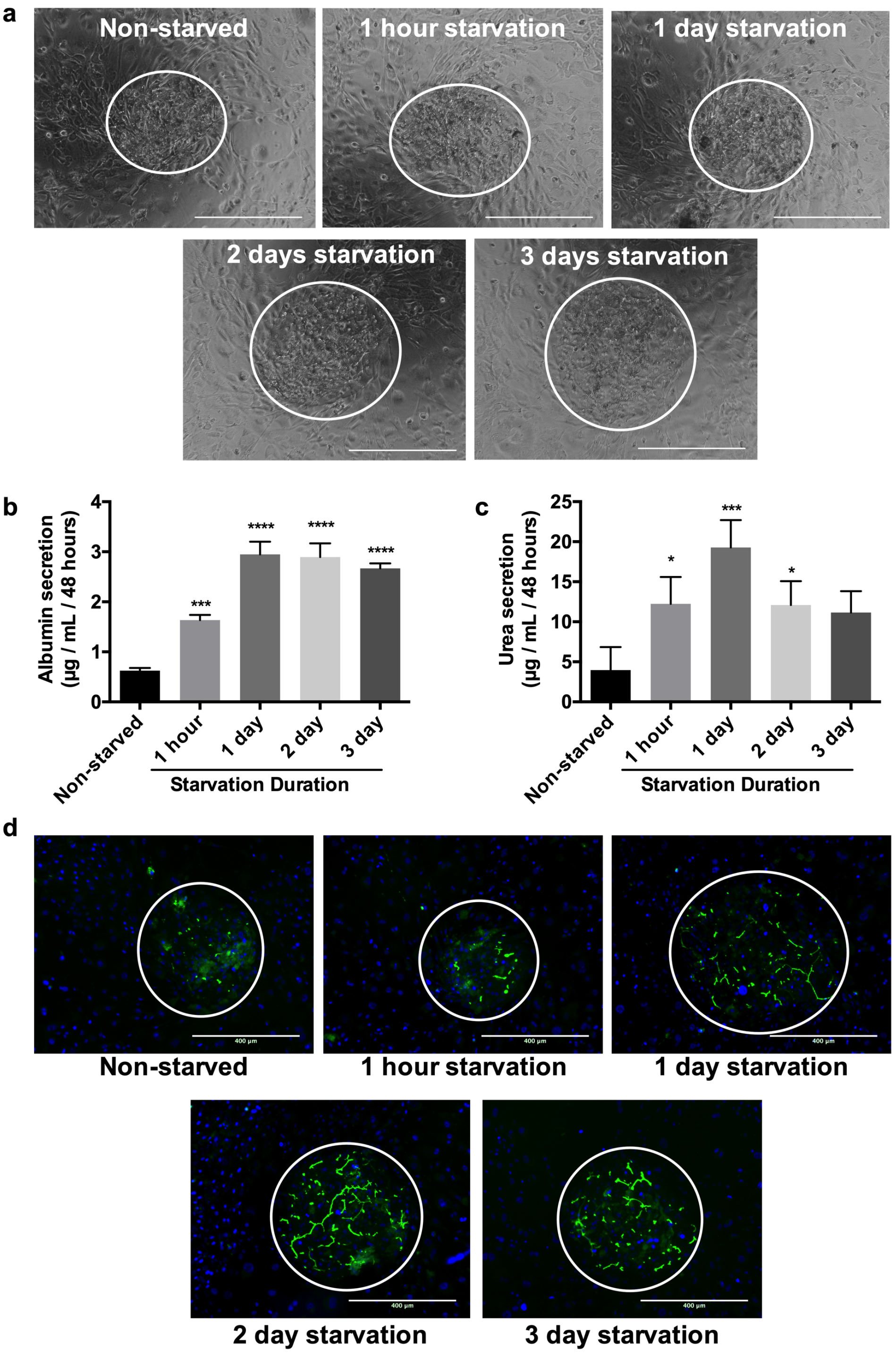


**Supplemental Figure 1. Optimization of intermittent starvation duration.** Micropatterned co-cultures (MPCC) containing primary human hepatocytes (PHH) and 3T3-J2 murine embryonic fibroblasts were created as described in Figure 1a of the main manuscript and cultured for 2 weeks in serum/hormone-supplemented maintenance culture medium. Then, MPCCs were washed with phosphate buffered saline and subjected to serum/hormone-free culture medium (starvation) for 1-hour, 1-day, 2-days, or 3-days, after which the cultures were put back in the serum/hormone-supplemented culture medium; such a starvation protocol was repeated weekly for 4 weeks. Non-starved MPCCs served as controls. (a) Phase contrast images of 6-week-old MPCCs subjected to different starvation durations as above and non-starved controls. (b) Albumin and (c) urea secretions from 6-week-old MPCCs subjected to different starvation durations as above and non-starved controls. (d) Functional bile canaliculi in PHHs within starved and non-starved MPCCs as assessed by the excretion of the CDF dye. Scale bars represent 400 µm. *$p<0.05$, ***$p \leq 0.001$, and ****$p \leq 0.0001$ relative to the non-starved control.

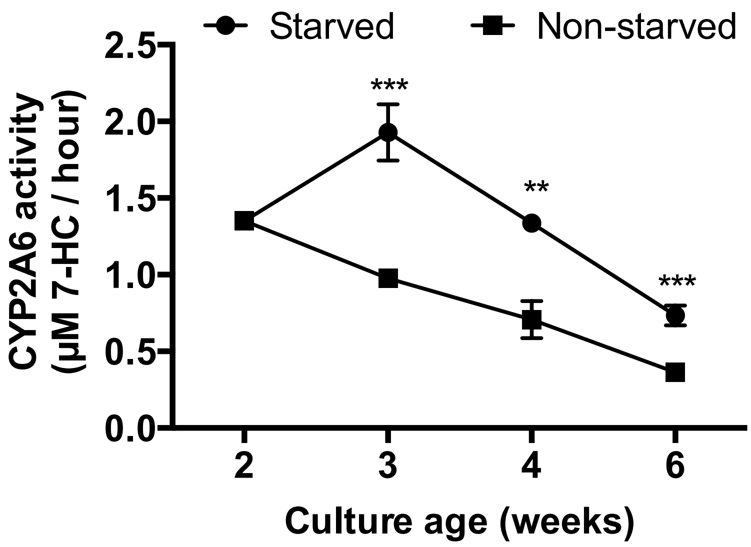


**Supplemental Figure 2. Intermittent starvation leads to higher CYP2A6 enzyme activity in hepatocytes.** MPCCs were starved for 2 days every week for 4 weeks as described in Supplemental Figure 1; non-starved MPCCs served as controls. CYP2A6 activity was quantified using the coumarin 7-hydroxylation reaction as described in the methods of the main manuscript. **$p<0.01$ and ***$p \leq 0.001$.

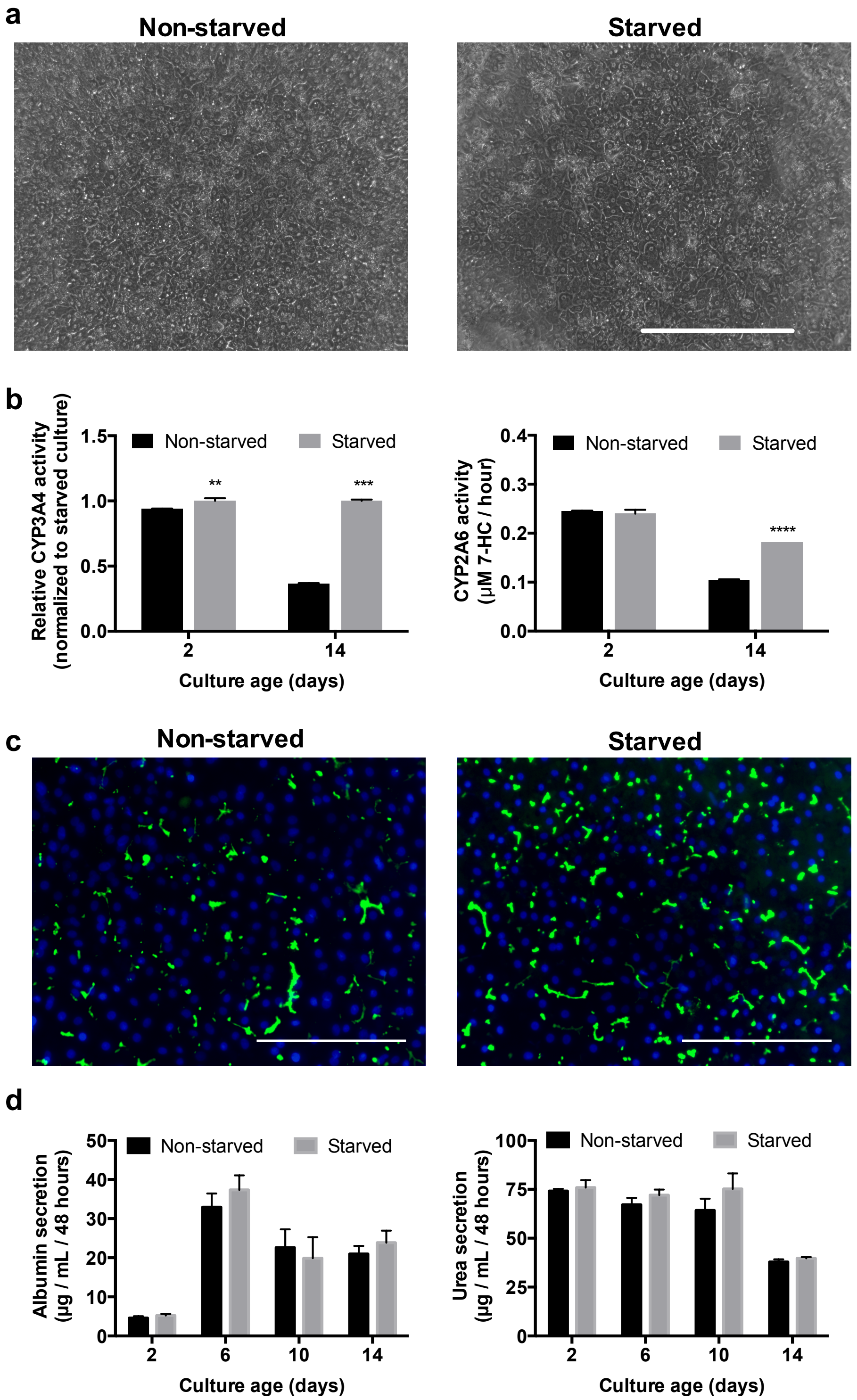


**Supplemental Figure 3. Effects of intermittent starvation on hepatocyte monocultures.** Cultures were periodically starved every two days for 2 weeks and assessed for (a) morphology, (b) CYP activities, (c) functionality of bile canaliculi via CDF secretion, and (d) secretions of albumin and urea in supernatants. Scale bars represents 400 µm. **p<0.01, ***$p \leq 0.001$, and ****$p \leq 0.0001$ relative to the non-starved control.

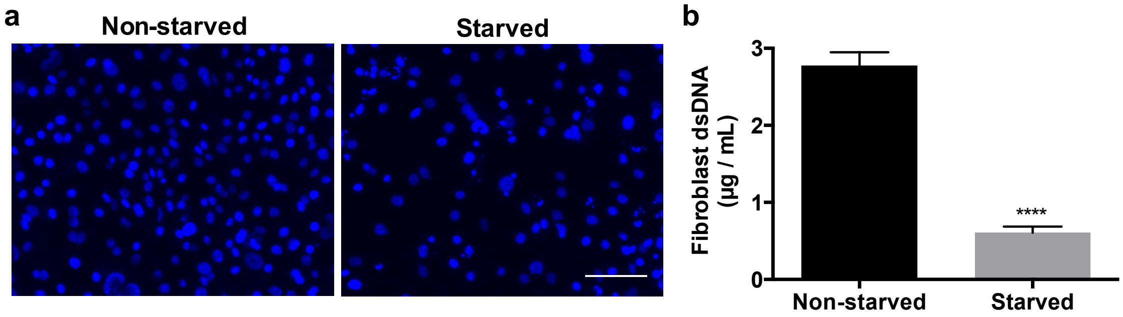


**Supplemental Figure 4. Intermittent serum starvation reduces fibroblast density.** (a) DAPI stained fixed fibroblast monocultures +/- 2-day serum starvation. (b) Fibroblast double stranded DNA (dsDNA) +/- 2-day serum starvation. Scale bar represents 100 µm. ****$p \leq 0.0001$.

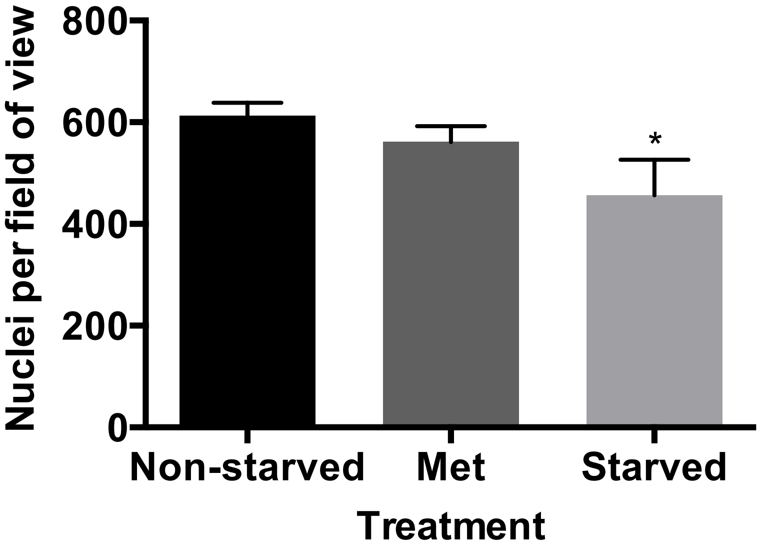


**Supplemental Figure 5. Effects of metformin treatment or intermittent starvation on fibroblast density in MPCCs.** MPCCs were starved for 2 days every week for 4 weeks as described in Supplemental Figure 1 or treated with metformin intermittently with the same timeline as starvation; non-starved MPCCs served as controls. Fibroblast density between PHH islands in 6-week-old MPCCs was assessed via live cell imaging of nuclei using Hoechst 33342 staining. *$p \leq 0.05$ relative to the non-starved control.

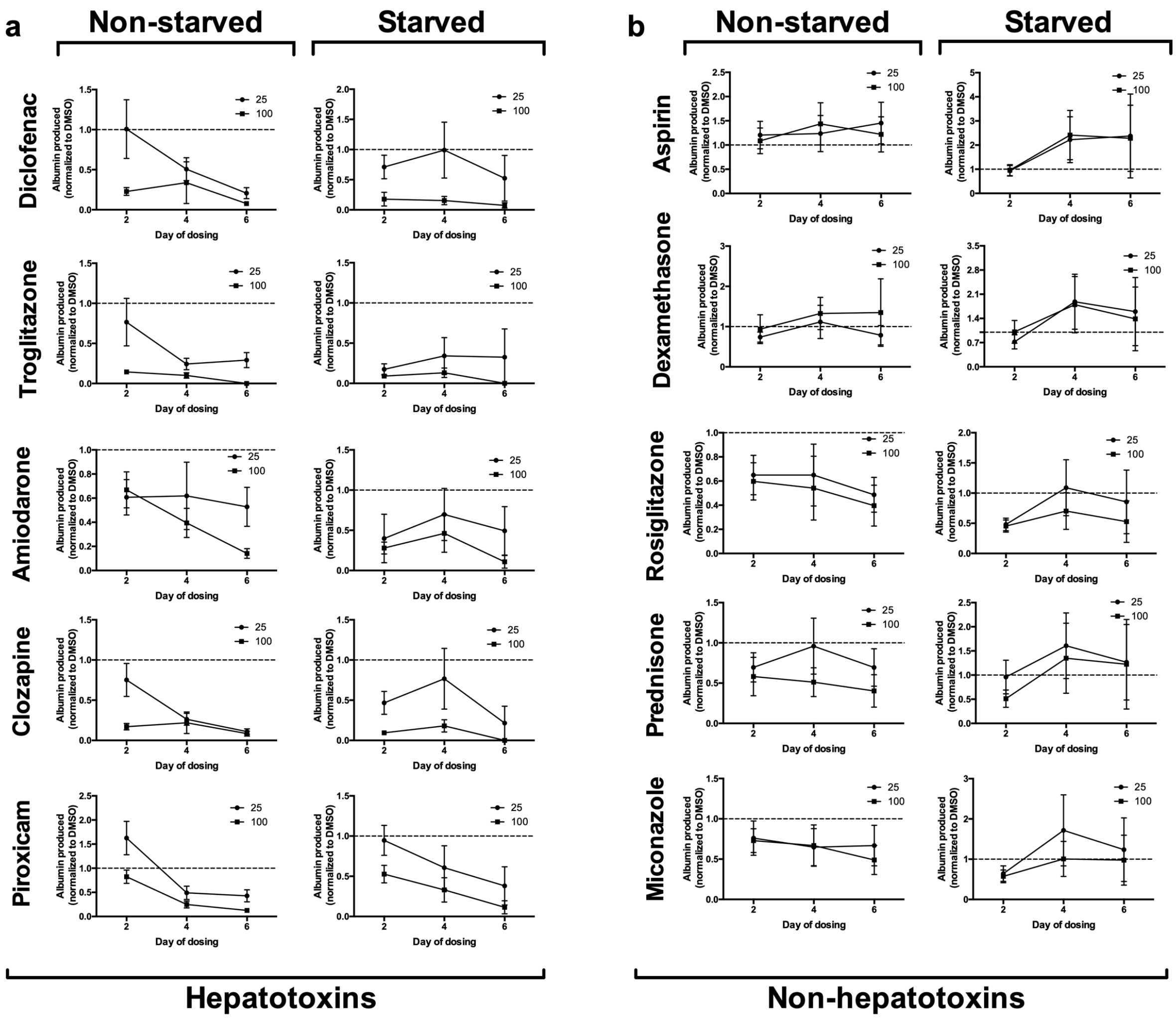


**Supplemental Figure 6. Albumin secretion in MPCCs treated with prototypical hepatotoxic and non-hepatotoxic drugs.** MPCCs were maintained either non-starved or starved for 2 days every week for 4 weeks and then treated with drugs for 6 days with prototypical hepatotoxins (left) and non-hepatotoxins (right) at 25× and 100× $C_{max}$ ($C_{max}$: maximum drug concentration measured in human plasma) for each drug with fresh drug added to culture medium at each 2-day medium exchange. Albumin secretion was assessed using an enzyme linked immunosorbent assay as described in the methods of the main manuscript. Data was normalized to the control cultures treated with DMSO alone.

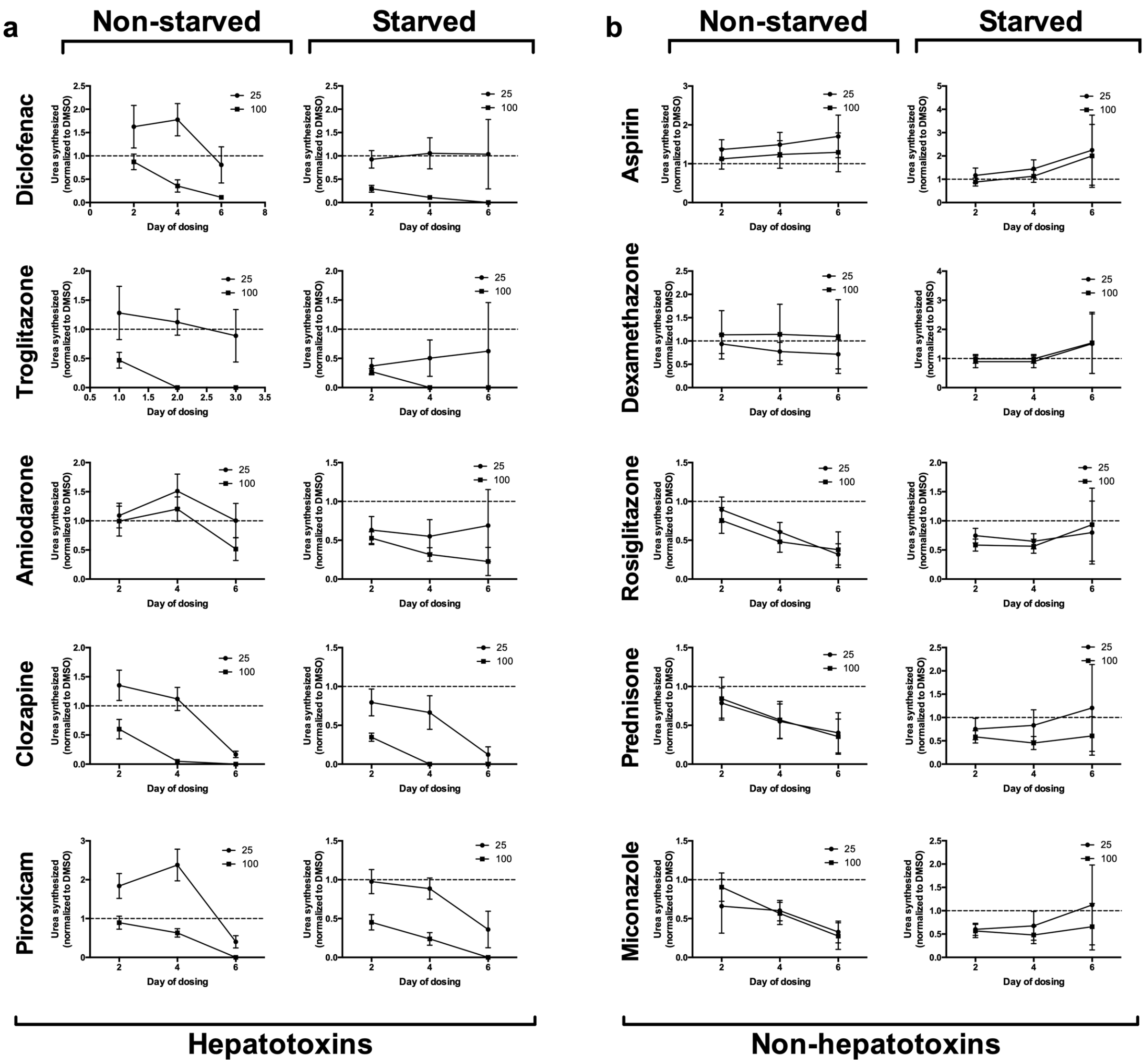


**Supplemental Figure 7. Urea secretion in MPCCs treated with prototypical hepatotoxic and non-hepatotoxic drugs.** MPCCs were created and treated with drugs as described in Supplemental Figure 6. Urea secretion was assessed using a colorimetric assay as described in the methods of the main manuscript. Data was normalized to the control cultures treated with DMSO alone.
